# Kinetics and targeting of Vipp1 aggregation in cyanobacteria

**DOI:** 10.1101/2022.12.01.518719

**Authors:** Colin Gates, Nicholas C. Hill, Kelsey Dahlgren, Jeffrey C. Cameron

## Abstract

The kinetics of regulation of vesicle-inducing protein in plastids (Vipp1), an essential protein in all oxygenic phototrophs, are not completely understood. Vipp1 is responsible for both membrane maintenance and damage repair processes, aggregating at sites of damage (proton leakage) or physical strain (responding to presence of anionic lipids). Herein we spatiotemporally resolve exchange between inactive cytoplasmic Vipp1 and these two functionalities. We demonstrate that Vipp1 can be aggregated to a specific site in the thylakoid membrane by laser damage, but this behavior appears to be mediated by a rapidly resolved secondary process, as cytoplasmic Vipp1 is rendered unresponsive to nearby damage on a time scale three orders of magnitude faster than the aggregation process itself.

**One Sentence Summary:** Vipp1 aggregation to points on the thylakoid is induced in response to specific damage and repair kinetics are observed.

## Main Text

Thylakoid membranes are the site of the initial steps of photosynthesis in oxygenic phototrophs across the tree of life. Accordingly, almost all molecular oxygen on Earth and a comparable fraction of the usable electrons in the biosphere were derived from water at a thylakoid membrane. The vesicle-inducing protein in plastids 1 (Vipp1, also known as inner membrane-associated protein of 30 kDa or IM30) is fundamentally associated with these membranes and similarly distributed throughout almost all photosynthetic organisms (*1*). Vipp1 is a dynamin-like protein closely related to the phage shock protein A (PspA) found widely in bacteria, though possessing a C-terminal domain necessary for thylakoid targeting which PspA does not have (*2*). It is essential for autotrophic growth in phototrophs across the tree of life and particularly in cyanobacteria, being necessary for thylakoid development and function (*3*). As a multifarious protein, Vipp1 has been found to be involved in a wide range of specific processes related to membrane-bound complex operation, assembly, and stoichiometry, structural reorganization of thylakoids, membrane biogenesis, repairs, and lipidome regulation (*4*). Complete depletion of Vipp1 has never been achieved under autotrophic conditions and knockdown has proven extremely detrimental to the light-dependent reactions of photosynthesis in all attempts (*5*–*7*).

Recent studies have established several regulatory mechanisms on the action of cyanobacterial Vipp1. The presence of two characteristic distributions was established by Bryan *et al*.; under suboptimal growth light levels, diffuse Vipp1 was observed in the cytoplasm, whereas application of supersaturating light resulted in the formation of microscopically visible puncta at the thylakoid membrane (*8*). It was subsequently shown that transition between Vipp1 expression states was related to a change in cell photosynthetic behavior rather than simply being controlled directly by light level (*6*). It is thought that the diffuse cytoplasmic phenotype represents a collection of monomers or small oligomers, while puncta are large ring-shaped complexes which fundamentally associate with the membrane. Among membrane-associated multimers, further differentiation of functionality has been established. Some Vipp1 complexes, particularly those associated with the thylakoid convergence zone (in cyanobacteria, the thylakoid organizing center; TOC), initiate membrane fusion events in response to availability of Mg^2+^ (*9*). The incidence of membrane fusion events is increased in response to leakage of protons (*10*). Membrane-associated Vipp1 complexes also play a more transient and less site-localized role in regulation of osmotic pressure on the thylakoid membrane to avoid damage (*11*). These are defined as “membrane destabilization” (bulk rearrangement including the vesicle formation in the protein’s name) and “membrane protection” (not involving rearrangement or fusion) processes. Despite its role in membrane reorganization, Vipp1 does not have any predicted transmembrane domains, but contains an amphipathic C-terminal domain which facilitates lipid association (*12*). The overall complex forms around a key repeating motif in which three peptides form a pocket which binds both Mg^2+^ and a purine di-or triphosphate which can be hydrolyzed by this and higher oligomeric groups (*13*). This hydrolysis appears to be at least part of the mechanism of oligomerization and growth of the ring structure which interacts with the membrane. However, the precise kinetics of aggregation in native conditions have not been studied to this point. Herein, we demonstrate the behavior of membrane-associated Vipp1 complexes in living cyanobacteria across multigenerational growth cycles and in response to precisely controlled damage events.

To investigate Vipp1 expression *in vivo*, we generated a mutant strain of *Synechococcus* sp. PCC 7002 with superfolder green fluorescent protein (GFP)-labeled Vipp1 (Vipp1-GFP hereafter) inserted into the genome in addition to the essential native Vipp1. This insertion has no discernable effect on growth rate or overall cell health and characteristic Vipp1 expression can be observed (Figure 1, Movie S1). Vipp1-GFP cells were grown in liquid medium to log-phase and diluted onto agarose pads as described previously (*14*) for observation of phenotype during growth from the single cell to microcolony stage. As shown in Figure 1A, upon initial imaging (following mild light stress and transition from liquid to solid medium) minor GFP puncta at positions including but not limited to the TOC (dual polar foci) are visible in cells. These puncta dissipate before the first division as cells acclimate to their new growth conditions, leaving a uniform background level of Vipp1-GFP evenly dispersed throughout the cytoplasm (7.5 h timepoint). This level is generally maintained, with occasional temporary aggregation in small puncta similar to those observed initially, in healthy cells. In contrast to these transient foci, a membrane damage event in the lower left position of cell “c” occurs at 8.5 h, resulting in a considerably greater accumulation of Vipp1-GFP to this site. This aggregation persists even through several cell divisions and is still visible 7.5 hours later at the end of the growth course in cell “o” (Figure 1A and 1B). A further consequence of this polar recruitment is the severe depletion of Vipp1-GFP in adjacent sister cell “g” following cell division (Figure 1C and 1D). Recovery of normal Vipp1-GFP levels takes nearly a full generation, potentially leading to increased susceptibility to membrane damage during this time in the depleted cell lineage. During the growth of this microcolony, a clear dichotomy is established between two types of Vipp1-GFP aggregates. Small, transient Vipp1-GFP puncta appear to occur for the purpose of local membrane strain alleviation; when these form, a background level of Vipp1 remains present in the cytoplasm and they generally dissipate quickly. In contrast, the larger Vipp1-GFP aggregates, as in Figure 1C, are long-lived and their formation is accomplished by accumulating almost all Vipp1 in the cell to that site. These appear to be formed in response to significant thylakoid membrane damage as indicated by co-localized patches of high chlorophyll fluorescence often associated with thylakoid membrane invaginations (Figure 1E). Additionally, these aggregates are sometimes observed to extend into larger filaments that may exceed the length of a single cell and transect the division site (Figure 1F).

**Fig. 1.**
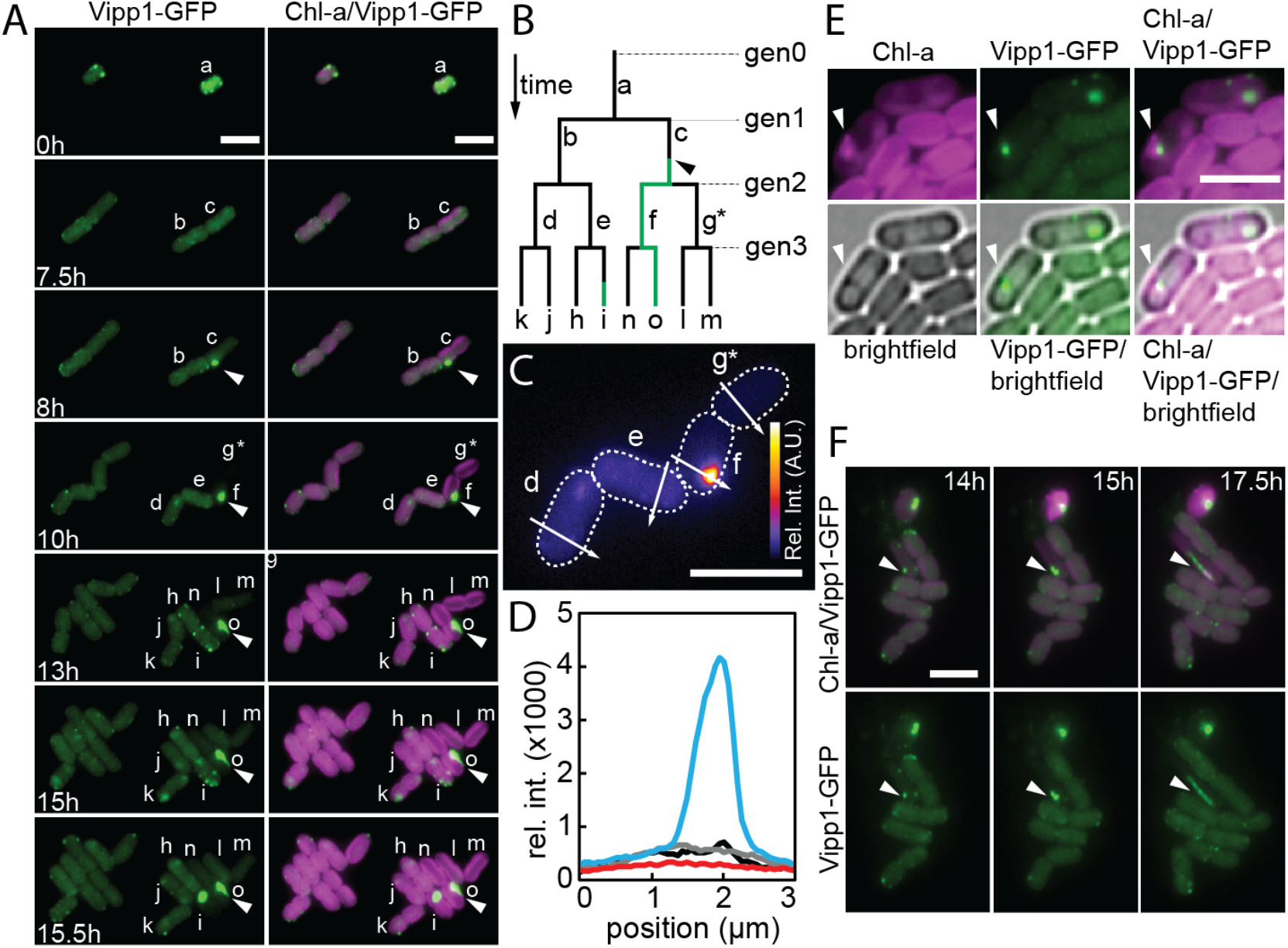
Multigenerational localization patterns of Vipp1-GFP in live cells during typical microcolony development. (A) Growth of two adjacent cells into microcolonies using time-lapse fluorescence microscopy. Aggregation of Vipp1-GFP during spontaneous damage event indicated by white arrow. Letters correspond to individual cell identities. (B) Schematic lineage tree with horizontal lines representing division events (generation indicated by gen#) and vertical lines representing individual cells. Labels correspond to cell identities shown in A. Vipp1-GFP aggregation events and multigenerational inheritance patterns indicated by green lines. Arrow indicates aggregation event in cell “c” highlighted in A. (C) Quantification of Vipp1-GFP aggregate in cell “f” and depletion of signal in sister cell “g*” following division of cell “c” compared to normal cells “d” and “e” (Calibration bar indicates relative fluorescent intensity of Vipp1-GFP between 150-4000 A.U.). (D) Line plot showing fluorescent intensity of Vipp1-GFP in panel C (solid arrows in panel C indicate position and direction of line scan). Cell “d”, black; cell “e”, grey; cell “f”, blue; cell “g*”, red. (E) Large Vipp1-GFP aggregate (white arrow; Green) in proximity to aberrant thylakoid membrane invagination with high fluorescence (magenta). (F) Time course showing filamentation of Vipp1-GFP aggregate. Chlorophyll-a (Chl-a; magenta) and Vipp-GFP (green) is shown in panels A, E, and F. All scalebars = 5 µm.

Using confocal microscopy and site-directed laser stimulation with sub-micron resolution, we investigated the damage necessary to induce Vipp1-GFP aggregation (Figure 2, Movie S3). While local stimulation of membrane-associated phycobilin (Figure 2A) with a 640 nm laser does not induce Vipp1 aggregation, targeting the Soret absorption of membrane-bound chlorophyll using a 405 nm laser induces an aggregation event of the damage type, consistent with what is observed during growth. It is probable that the spectral specificity is unrelated to chlorophyll altogether and is a simple membrane damage event which exposes certain lipids to reactive oxygen species (*15*), which may be the direct signal for Vipp1 aggregation. Immediately after laser stimulation in Figure 2B, Vipp1-GFP remains evenly distributed throughout the cytoplasm. Subsequently, after 6 minutes a significant Vipp1-GFP aggregate is visible at the 405 nm stimulation site. By ten minutes essentially all Vipp1-GFP in the cell is localized exclusively to the 405 nm damage site and no significant release occurs within 42 minutes (Figure 2C). As seen in Figure 2B and 2C, a consistent level of Vipp1-GFP is maintained in the cell during the repair process, none of which is present away from the damage site. A similar phenomenon is observed under native growth conditions when the damage event occurs shortly before cell division (Figure 1, Movie S1 and S2). Cell division continues at roughly the same rate as it would have, resulting in one daughter cell having essentially two cells’ Vipp1 present, while the other daughter cell has no Vipp1 and takes several hours to produce more. During this time, the damaged cell does not appear to generate any cytoplasmic Vipp1 of its own.

**Fig. 2.**
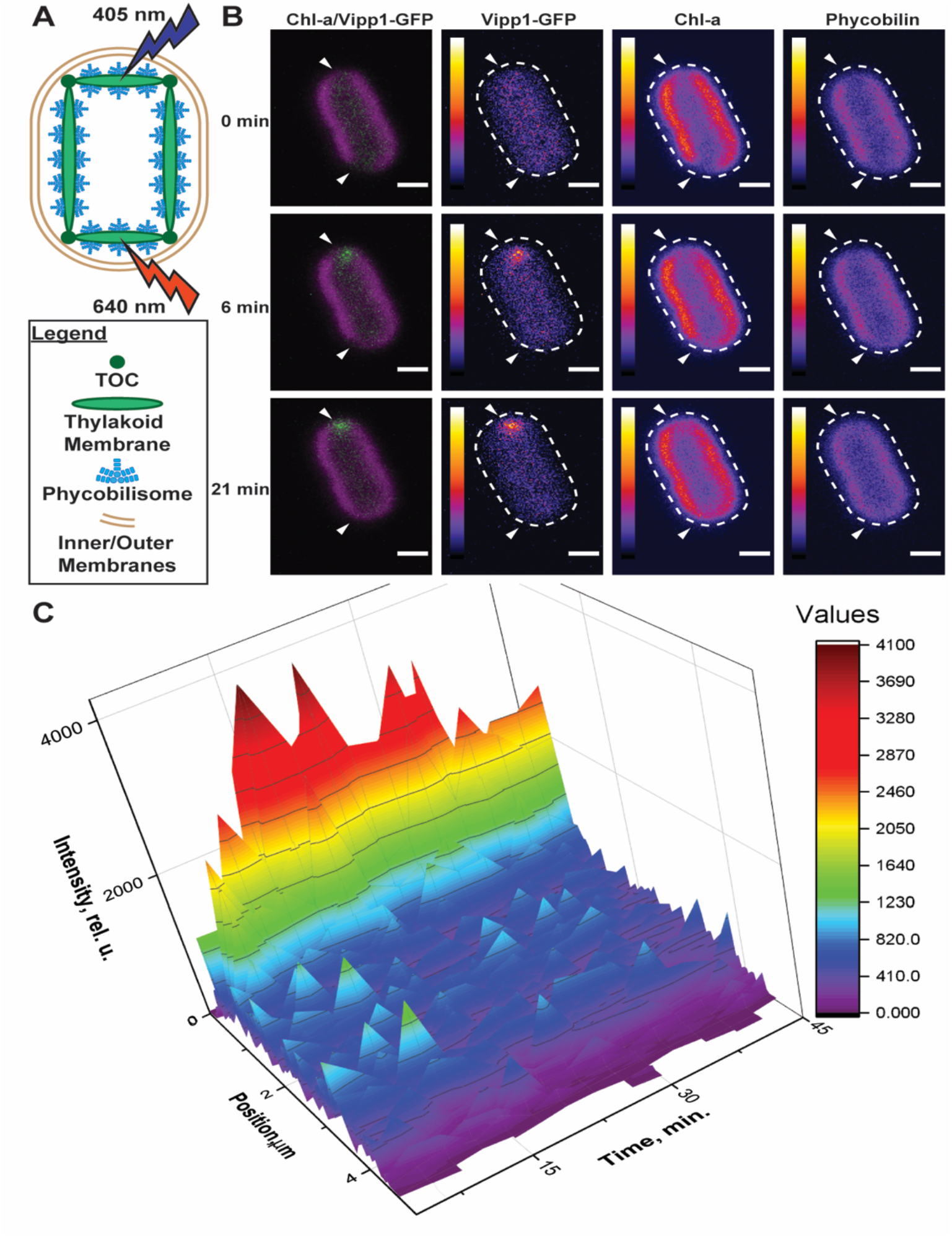
Spectral selectivity and kinetics of Vipp1 recruitment in a single cell. (A) Schematic diagram of cyanobacterial cell and experimental setup. (B) Time series of a representative cell imaged immediately after a 2 ms laser stimulation with 405 nm light at top of cell and 640 nm light at bottom of cell. White arrows indicate approximate position of laser stimulation. Scale bars = 1 µm. Calibration bars indicate relative fluorescent intensity between 0-4095 A.U. Beginning at ∼3 min, Vipp1-GFP aggregates exclusively to the site of 405 nm stimulation and does not aggregate at the 640 nm site over the duration of the experiment. Loss and recovery of pigment (phycobilin and Chl-a) fluorescence due to laser stimulation is similar between both sites. Still frames extracted from Movie S3. In the first column, Chl-a fluorescence is shown in magenta and Vipp1-GFP in green. (C) Three-dimensional plot of Vipp1-GFP fluorescence intensity along a line through the cell that transects both laser stimulation sites over 42 minutes.

While damage repair is ongoing, the cell has a limited ability to respond to subsequent membrane damage events at secondary sites as the Vipp-GFP aggregation process is set in motion much more quickly than the actual formation of the aggregate. In Figure 3 (Movie S4), two sites at opposite ends of a healthy cell are stimulated with a 405 nm laser 100 ms apart (Figure 3A). This is a gap three orders of magnitude smaller than the time required to fully form the aggregate observed in Figure 2. Yet, after 6 minutes, the site that was stimulated first has aggregated almost all free Vipp1-GFP in the cell while the second site has aggregated none above the background level (Figure 3B). As seen in Figure 3C, release of Vipp1-GFP at the initial site occurs concurrently with aggregation at the second site, although little Vipp-GFP is observed in the cytoplasm during this exchange. After 42 minutes (Figure 3C) the initial damage site is mostly repaired and the majority of Vipp1-GFP has aggregated at the second site, where repair is ongoing.

**Fig. 3.**
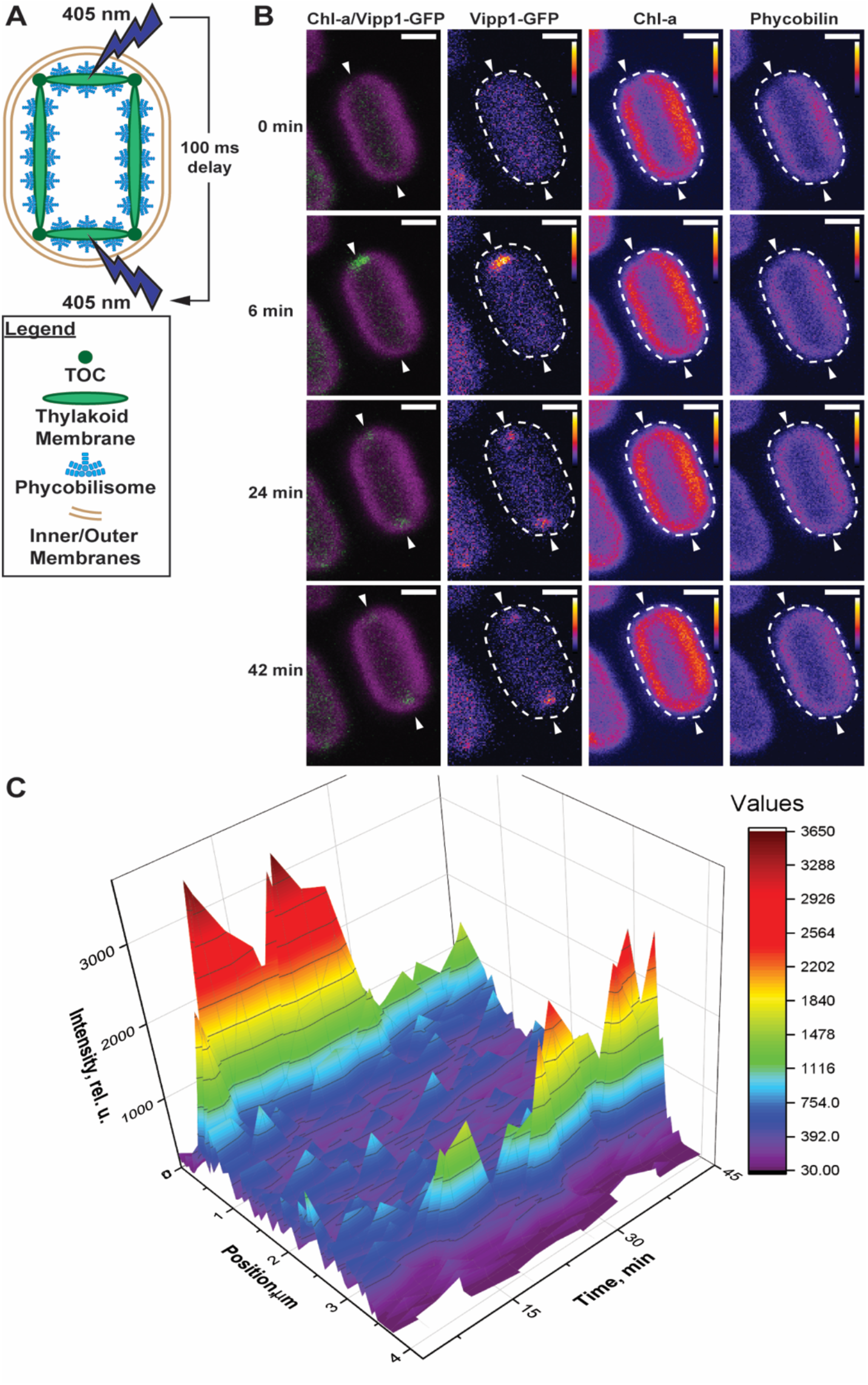
Kinetics and localization of Vipp-GFP recruitment in a single cell following stimulation with consecutive 405 nm laser pulses. (A) Schematic diagram of cyanobacterial cell and experimental setup. (B) A representative cell imaged immediately after stimulation with a 2 ms laser pulse of 405 nm light at both ends. Aggregation of Vipp1-GFP is first observed exclusively at the first (top) site by ∼6 minutes. By 24 minutes, Vipp1-GFP is observed at both sites. By 42 minutes, almost all Vipp1-GFP is concentrated at the second site (bottom) and the pigment (Chl-a and phycobilin) fluorescence at the top site is fully restored. Still frames were extracted from Movie S4. In the first column, chlorophyll-a fluorescence is shown in magenta and Vipp1-GFP in green. Scale bars = 1 µm. (C) Three-dimensional plot of Vipp1-GFP fluorescence intensity along a line through the cell that transects both sites of laser stimulation over the 42-minute time course.

The enormous disparity between the time required to target the entirety of a cell’s Vipp1-GFP to a particular damage site and the actual aggregation of the oligomeric structure at that site suggests a secondary, and potentially unexplored, regulatory mechanism. Otherwise, at least two of the necessary factors (Vipp1 monomers and a locally acidified region (*10*)) are present at the site of secondary damage in Figure 3 early enough that some Vipp1-GFP should be recruited to this site. Nevertheless, oligomerization at the second site does not occur until a fraction of Vipp1-GFP is lost from the initial site of damage. This suggests that these factors alone are inadequate to regulate Vipp1 oligomerization and the other factors, Mg^2+^ and ATP/GTP (*13*), may not be present. The N-terminal alpha helix, previously posited as a detector for membrane stress at even the monomeric level (*16*), similarly does not appear to be the primary determinant of oligomer localization in damage conditions, as once again there is Vipp1-GFP in close proximity to the second damage site long after damage occurs, but this Vipp1-GFP exclusively oligomerizes at the first damage site in a process akin to Ostwald ripening. It is possible that the recruitment of Vipp1-GFP to a particular site is triggered by the presence and activity of a related but faster-acting dynamin-like protein such as DynA, which serves as a temporary seal on the membrane in response to pore formation and is also regulated by low pH and GTP availability (*17, 18*). Critically, DynA does not permanently reseal the membrane and its presence causes localization of the type of anionic lipids known to recruit Vipp1. DynA has previously been proposed as a candidate for triggering hole repair (*17*), and a protein of this type could serve as a more effective signal for Vipp1 oligomerization. Given the distribution of dynamin-like proteins across the tree of life (*19*), these recruitment kinetics may play a role in other critical settings. In addition to PspA, the eukaryotic ESCRT-III has a similar structure and function (*20*) and may follow similar behavior patterns. Mammalian viral response proteins, such as the myxovirus and HIV response protein MxA (*21*), may also follow similar response kinetics for direct membrane damage. This method for directed production of minor punctures in the membrane will allow further analysis of cellular Vipp1 activity and oligomerization kinetics and shows considerable promise for similar studies in other systems in which their membranes can be perforated by targeted laser stimulation.

## Author contributions

Conceptualization: J.C.C. and C.G. Methodology: all authors. Analysis: C.G. and J.C.C. Investigation: C.G., N.C.H., and K.D. Writing: C.G. and J.C.C. (draft, figures) and all authors (final draft);

## Data and materials availability

All data is available in the main text or the supplementary materials.

## Supplementary Materials

Materials and Methods

Movies S1-S4

References (*21-24*)

## Supplementary Materials

## Materials and Methods

### Construction of the Vipp1-GFP strain

The *Vipp1* gene (SynPCC7002_A0294) was amplified from PCC 7002. Plasmids were assembled using Gibson Assembly (*22*) with superfolder GFP (*23*) as a C-terminal fusion on Vipp1, NS1 as the homology arms, p_*ccmK2*_ as the promoter (*24*), and kanamycin resistance for selection. The Gibson reactions were transformed into DH5α *E. coli*, and minipreps of liquid cultures started from single colonies were performed to collect plasmid. This plasmid was transformed into PCC 7002 and colonies containing the desired insert were serially passaged under growth conditions as given below, in the presence of 100 μg/ml kanamycin, until segregated.

### Cultivation

PCC 7002 cultures were grown in an AL-41 L4 Environmental Chamber (Percival Scientific, Perry, IA) at 37°C under 150 μmol photons m^-2^ s^-1^ constant illumination from cool white fluorescent lamps under air (0.04% CO_2_). Cultures were grown in 25 ml of A^+^ media (*25*) with orbital shaking at 100 rpm in 125 ml baffled flasks with foam stoppers, or on 1% w/v agar plates of pH 8.2 A^+^ media. All growth media was supplemented with 100 μg/ml kanamycin to retain the pure mutant line.

### Microscopy

For long-term imaging (Figure 1, Movie S1), all images were obtained on a Nikon TiE custom inverted wide-field microscope with Perfect Focus System. Temperature in all images and the growth series was maintained at 37°C using a Lexan environmental chamber (BioSpherix, Parish, NY). Trans-illuminating growth light was supplied from a light-emitting diode (LED) light source (Lida Light Engine, Lumencor, Beaverton, OR). Epifluorescence imaging light was supplied from a custom-filtered LED light source (Spectra X Light Engine, Lumencor, Beaverton, OR) and delivery was controlled using a synchronized hardware-triggered shutter. The microscope was controlled using NIS Elements AR software (version 5.11.00 64-bits) with Jobs acquisition upgrade. Images were acquired using an ORCA-Flash4.0 V2+ Digital sCMOS camera (Hamamatsu) with a Nikon CF160 Plan Apochromat Lambda 100× oil immersion objective (1.45 numerical aperture). Cells imaged were obtained from liquid culture in exponential phase (*24*) and diluted to 0.05 OD_730 nm_, from which 2 μl was spotted onto a 1% agarose A^+^ pad (with 100 μg/ml kanamycin) which had been incubated at 37°C for 1 hour. Cells were dried onto the pad and inverted onto a 35-mm imaging dish (ibidi μ-dish) with glass coverslip bottom, which was wrapped with Parafilm M to prevent drying of the pad. Cells were grown under 37°C and 150 μmol photons m^-2^ s^-1^ red light centered at 640 nm; this light was maintained at all times except during fluorescent imaging. Images were taken every 30 minutes using 470- and 640-nm excitation from the Spectra X light engine and emission collected using standard GFP (green in images) and Cy5 (magenta in images) filters (Nikon).

For short-term imaging (Figure 2, 3, Movie S3, S4), all images were obtained on a customized Olympus FV-3000 confocal microscopy system with Olympus UPlanXApo 60x oil immersion objective (1.42 numerical aperture) built onto an Olympus IX-83 inverted fluorescence microscope with Z-Drift Compensator and onboard excitation and imaging illumination from lasers at 405, 488, 561, and 640 nm (50, 50, 50, and 40 mW Coherent OBIS laser modules, respectively). Cells were streaked onto agar pads and inverted onto 4-well ibidi μ-Slides and allowed to acclimate to imaging conditions on the microscope following the observation that Vipp1-GFP is delocalized in acclimated cells (Figure 1). “Phycobilin” measurements (blue) were performed using an excitation laser power of 0.02%, wavelength of 640 nm, and emission range of 650-720 nm. This wavelength excites both allophycocyanin (primary absorption) and chlorophyll and thus some of the yield here is from chlorophyll as well. “Chlorophyll-a” measurements (magenta) were performed using an excitation laser power of 0.05%, wavelength of 405 nm (which does not also excite phycobilins), and emission range of 650-720 nm. GFP measurements (green) were performed with standard EGFP settings (Olympus) of 488 nm excitation and 510-540 nm emission range with laser power 0.05%. All imaging was performed at a capture time of 2.0 μs/pixel. Individual cells without GFP puncta (Vipp1-GFP aggregation sites) were stimulated using 405 or 640 nm laser at 10% power applied for 2 ms (20,000x imaging light intensity at 405 nm and 50,000x imaging light intensity at 640 nm) at a specific site within the cell using Olympus FV31S-SW LSM Stimulation, after which Olympus Sequence Manager was used to image the cell at 3-minute intervals for a total of 42 minutes using phycobilin, chlorophyll-a, and GFP channels.

### Image Analysis

To quantify spatiotemporal resolution of GFP localization in Figures 2 and 3 and Movies S3 and S4, pixel intensity over a line plot through bleach sites was obtained for each image using ImageJ (Fiji). These line intensity plots were then plotted by frame over time as a 3D surface using OriginLab Origin 2018b. Spatiotemporal resolution of GFP expression from Movie S1 was extracted into Movie S2 using a surface plot of the GFP channel over the entire movie using a “Fire” lookup table (Fiji).

---

**Movie S1**.

Stages of microcolony growth, showing characteristic microcolony morphology in *Synechococcus* sp. PCC 7002, and natural Vipp1-GFP localization during this time. Chlorophyll-a (Chl-a) fluorescence in magenta, Vipp1-GFP fluorescence in green. Frames are 30 minutes apart. Scale bar = 5 µm.

**Movie S2**.

Spatial resolution of Vipp1-GFP localization in the microcolony which suffers damage events in Movie S1 and Figure 1. Vipp1-GFP is shown using a “Fire” lookup table (ImageJ) in which higher intensity appears red and lower intensity appears blue. Frames are 30 minutes apart.

**Movie S3**.

Kinetics and damage-specificity of Vipp1 recruitment in a single cell following stimulation of points in the cell with 2 ms exposure to laser light. Top site: 405 nm light; bottom site: 640 nm light. Phycobilin fluorescence in red, chlorophyll fluorescence in blue, Vipp1-GFP fluorescence in green. Frames are 3 minutes apart.

**Movie S4**.

Kinetics and damage-specificity of Vipp1 recruitment in a single cell following stimulation of points in the cell with 2 ms exposure to 405 nm laser light and 100 ms delay between stimulation of top site and bottom site. Phycobilin fluorescence in red, chlorophyll fluorescence in blue, Vipp1-GFP fluorescence in green. Frames are 3 minutes apart.

